# Influence of endosymbionts on the reproductive fitness of the tick *Ornithodoros moubata*

**DOI:** 10.1101/2023.05.09.539061

**Authors:** Taraveau Florian, Pollet Thomas, Duhayon Maxime, Gardès Laëtitia, Jourdan-Pineau Hélène

## Abstract

Over the past decade, many studies have demonstrated the crucial role of the tick microbiome in tick biology. The soft tick *Ornithodoros moubata* is a hematophagous ectoparasite of *Suidae*, best known for transmitting the *African swine fever virus*. Its bacterial microbiota is characterized by a high prevalence of *Francisella*-like and *Rickettsia* endosymbionts. The present study aims to better understand the potential influence of these two major members of the tick microbiota on the reproductive fitness of *O. moubata*. A total of 132 adult female ticks were treated with antibiotics using gentamycin or rifampicin added to the blood meal. Half of the ticks were also supplemented with B vitamins to address the nutritional role of endosymbionts. Over two periods of 50 days, several traits related to the reproductive fitness were monitored to investigate the importance of *Francisella* and *Rickettsia* for these traits. It appeared that most of the reproductive parameters considered were not affected. However, antibiotic treatments induced an increase in the tick survival, indicating a potential fitness cost of harboring endosymbionts during the tick reproductive cycle. Similarly, 366 first stage nymphs of *Ornithodoros moubata* were exposed to the same treatments for molecular quantification of both endosymbionts. Results from qPCR suggested that the treatments had a bacteriostatic effect on endosymbionts without completely eliminating neither *Francisella*-like endosymbiont nor *Rickettsia*.

## Introduction

While ticks are among the most important vectors of pathogens affecting humans and animals worldwide, it is now well established that tick-borne pathogens (TBPs) coexist with many other microorganisms in ticks (e.g., Bonnet *et al*. 2017; Estrada-Peña *et al*. 2018). These other non pathogenic tick-borne microbes (microbiota) are likely to confer multiple detrimental, neutral, or beneficial effects to their tick hosts (Duron *et al*. 2018; Duron, Cremaschi & McCoy 2016). In addition, many studies have identified multiple interactions between some members of the tick microbiota and TBPs (e.g., Lejal *et al*. 2021; Aivelo, Norberg, & Tschirren 2019). For instance, these tick microbial communities may contribute to the transmission or multiplication of TBPs (Vayssier-Taussat *et al*. 2014; Bonnet *et al*. 2017; Budachetri *et al*. 2018; Bonnet & Pollet 2021), raising the question of whether tick microbiomes can be manipulated and used as a new tool in tick and tick-borne disease control.

Besides tick-borne pathogens, the other tick microbes can be broadly classified as endosymbiotic or commensal. Endosymbionts are intracellular bacteria, maternally transmitted and categorized as primary or secondary endosymbionts according to the tick species and their importance for the host (Jiménez-Cortés *et al*. 2018). Primary symbionts are obligatory, inheritable symbionts with a crucial role in the tick survival and development (Duron *et al*., 2018; Bonnet *et al*. 2017). In contrast, secondary endosymbionts are facultative and not required for host survival. They are known to manipulate reproduction, protect ticks against natural enemies or facilitate adaptation to changing environments (Duron *et al*., 2018). Finally, commensal bacteria are acquired from the environment and may differ from one individual to another. They are not transmitted vertically, their biological effects on ticks remain largely unknown, even though some recent studies suggest that some of them may participate in blood meal digestion (Narasimhan & Fikrig 2015) or influence pathogen presence or transmission (e.g., Adegoke *et al*. 2020; Lejal *et al*. 2021).

Four main bacterial genera are currently well identified as tick primary endosymbionts: *Coxiella-*like endosymbionts (CLE), *Candidatus Midichloria mitochondrii, Francisella-*like endosymbionts (FLE), and, less often, *Rickettsia*. All of these endosymbionts were identified to carry a copy of genes involved in essential nutrients synthesis, including B vitamins which are not naturally brought by the strictly hematophagous nutrition. Depending on the tick species, one or two of these genera will be found predominantly and will assume the nutritional function (Duron & Gottlieb 2020; Bonnet & Pollet 2021). For instance, this role is assumed by *Coxiella*-like endosymbionts (CLE) in *Amblyomma americanum* whereas it is assumed by *Rickettsia* in *Ixodes pacificus* (Rio, Attardo & Weiss 2016). Over time, several evolutionary events of loss and/or new endosymbiosis are suspected, leading to the diversity of endosymbionts currently found (Duron *et al*. 2017; Duron & Gottlieb 2020). However, the role of primary endosymbionts is not limited to nutrient supply. In ticks, they are also known to have an influence on fitness, reproduction (Guizzo *et al*. 2017; Li, Zhang, & Zhu 2018; Zhang *et al*. 2017; Zhong, Jasinskas, & Barbour 2007; Kurlovs *et al*. 2014), immunity, and vector competence (Narasimhan *et al*. 2014; Bonnet & Pollet 2021; Gall *et al*. 2016).

The life cycle of soft ticks presents specific characteristics not found in hard ticks: several nymphal stages, brief blood meals, adult females feeding and laying eggs multiple times before dying. These characteristics make soft ticks an ideal model for studying, in laboratory, the role of the tick microbiota on tick biology and physiology. *Ornithodoros moubata* is a soft tick widely distributed in Southern and Eastern Africa (Vial 2009; Leeson 1952), where its main hosts include wild and domestic members of the *Suidae* family: warthog, bushpig and domestic pig (Peirce 1974). This tick species is mainly known as a reservoir host for the *African swine fever virus* (ASFV) and is consequently an important concern for animal health issues (Sánchez-Vizcaíno *et al*. 2015). High-throughput 16S rDNA sequencing in *O. moubata* revealed that the most abundant bacterial species belonged to the genus *Francisella* and *Rickettsia* (Duron *et al*. 2018; Piloto-Sardiñas *et al*. 2023). Both bacteria are recognized as maternally-inherited endosymbionts in ticks (Duron *et al*. 2017). Furthermore, *Francisella*-like endosymbionts (FLE) have been shown to play a key role in the synthesis of several B vitamins, including biotin, folic acid and riboflavin, which are deficient in the blood meal (Duron *et al*. 2018).

Apart from their role in tick nutrition, the influence of *Francisella* and *Rickettsia* in the physiology and life history traits of soft ticks remains poorly known. In comparison, past studies suggest a role of CLE and *Rickettsia* for the reproductive fitness of female hard ticks belonging to different genera (Kurlovs *et al*. 2014; Li, Zhang, & Zhu 2018; Zhang *et al*. 2017; Zhong, Jasinskas, & Barbour 2007). In these studies, the reproductive fitness (number of eggs laid, mass of eggs, duration of oviposition and hatching rate) was negatively impacted by the elimination of parts of the microbiota.

The present study aimed to better characterize the role of key members of the microbiota in reproductive fitness in soft ticks. Accordingly, our first objective was to eliminate with antibiotic treatments the main endosymbionts (namely FLE and Rickettsia) of the microbiota of *Ornithodoros moubata*. The second goal was to evaluate the consequences of such disruption on the reproductive fitness of ticks, using vitamin B supplementation to identify whether part of the observed changes was directly attributable to an action of the endosymbionts on the reproduction or if it was a consequence of their nutritional role.

## Material and methods

### Ticks

All ticks were issued from a colony maintained at the CIRAD laboratory (Montpellier, France, member of the Vectopole Sud network) since 2008. This colony originated from the Neuchâtel strain initially collected in Tanzania and maintained at the University of Neuchâtel (Switzerland).

The female ticks used for the experiments had never been mated before and did not receive any new blood meal since molting from the last nymphal stage to the adult stage.

Adult ticks, nymphs, larvae, and eggs were kept in stable conditions in one incubator, at 25°C, 85% relative humidity, and 0% luminosity.

### Overview of the experimental design

Two experiments were conducted independently (**Figure 1**). In experiment one, first stage nymphs were treated once with antibiotics and B vitamins, and collected at different times for DNA extraction and qPCR targeting FLE and *Rickettsia*. In experiment two, adult females were treated with antibiotics and B vitamins in two successive treatments, and traits associated with reproduction were monitored after each treatment. At the end of this second experiment, females were collected for DNA extraction and qPCR targeting FLE and *Rickettsia*. The ticks used in these experiments were not offspring of one another.

**Figure 1:**
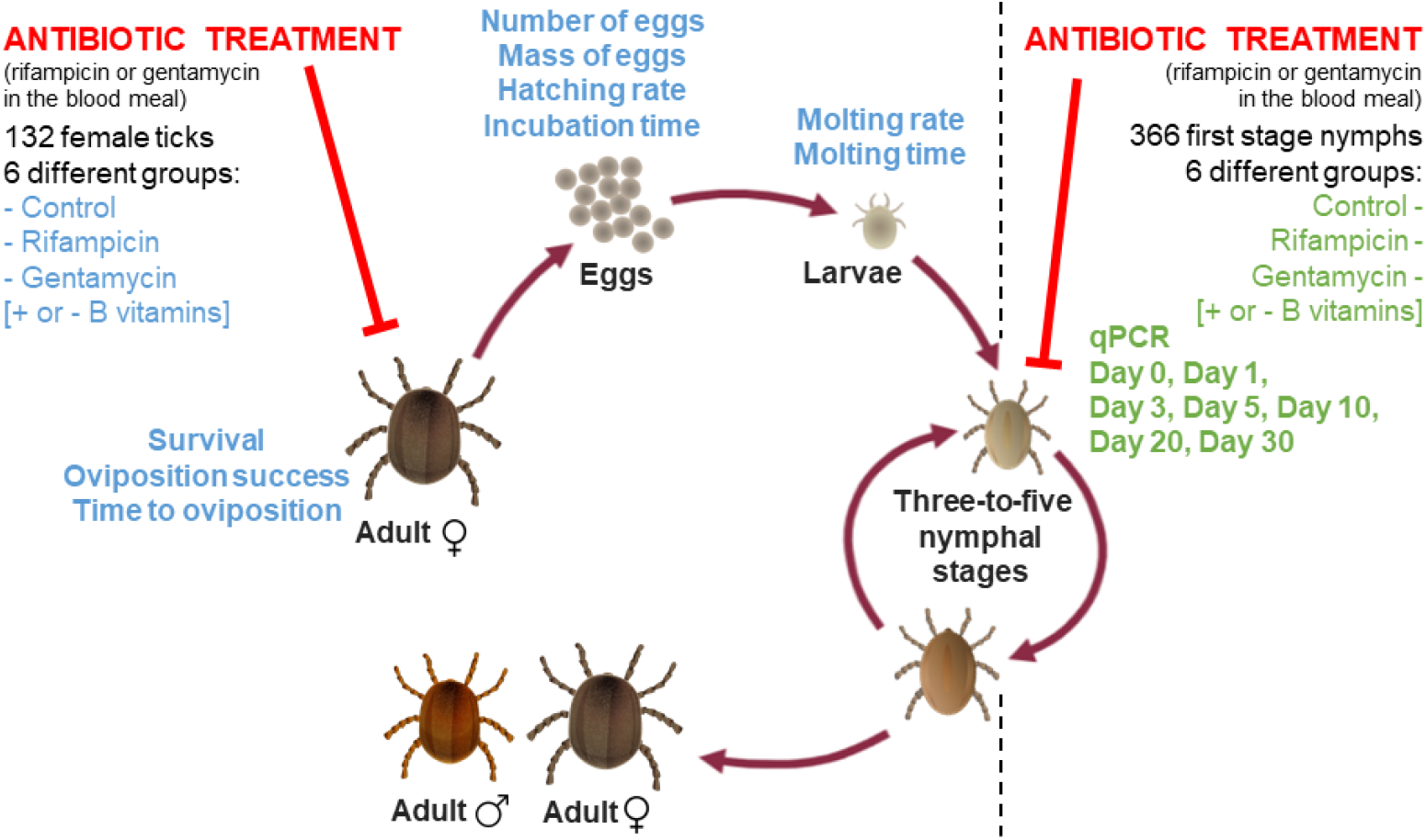
Experimental design for the use of **antibiotic treatments, followed by the evaluation of traits of interest at different stages of the parasitic cycle of *Ornithodoros moubata***. The traits monitored are indicated near the concerned parasitic stage. In green, the experiment 1 conducted on first stage nymphs to monitor bacterial loads after antibiotic treatments. In blue, the experiment 2 conducted on adult ticks to monitor reproductive parameters. Experiment 1 and 2 were performed independently from one another. Tick pictures: Françoise Chirara © Cirad-ASTRE.

Six treatment combinations were used in both experiments: Control (no antibiotics) without B vitamins, Control with B vitamins, Gentamycin without B vitamins, Gentamycin with B vitamins, Rifampicin without B vitamins, and Rifampicin with B vitamins.

### Feeding and antibiotic treatments

Antibiotic treatments and B vitamins were added to the blood used for blood feeding. First stage nymphs were engorged in groups of thirty individuals for one treated blood meal. Bloodmeal was performed using an artificial feeding system: culture plates with fresh cow blood inside the wells and a paraffin membrane above it for ticks to bite through. Blood was collected 1-3 hours before the experiment in 10mL heparinized tubes and all the blood used for this experiment was obtained from the same cow. Blood meals lasted three hours in complete darkness while blood temperature was maintained at 41°C and blood was stirred at 250 rpm. For each blood meal, a 1mL blood sample was collected for DNA extraction and qPCR targeting FLE and *Rickettsia*. Antibiotics and B vitamins were used following a modified protocol (Duron *et al*. 2018) with dilutions performed in Milli-Q water instead of methanol due to the potential toxicity of methanol for arthropods (Kaviraj, Bhunia & Saha 2004). Concentrations were chosen according to recommendations for sensitive bacterial species and previous studies in the literature: gentamycin 10 μg.mL^-1^, rifampicin 10 μg.mL^-1^, biotin 1 μg.mL^-1^, folic acid 30 μg.mL^-1^, riboflavin 20 μg.mL^-1^ (Duron *et al*. 2018; Zhong, Jasinskas & Barbour 2007; Narasimhan *et al*. 2014). Antibiotics and B vitamins were replaced by the same volumes of Milli-Q water for negative controls.

The choice of antibiotic molecules was based on the targeted endosymbionts, namely FLE and *Rickettsia* (Duron *et al*. 2018). Based on this study, it was decided to use rifampicin which can eliminate both Rickettsia and FLE, and gentamycin, for which most *Rickettsia* strains are resistant whereas *Francisella* species are sensitive (Vanrompay *et al*. 2018). Therefore, the elimination of *Francisella* alone under gentamycin treatment would allow to evaluate its role independently from *Rickettsia* (for which insights would be given using rifampicin treatment). Other molecules such as kanamycin and tetracycline were ruled out due to their potential toxicity for arthropods (Li, Zhang & Zhu 2018; Coon, Brown & Strand 2016). Finally, the vitamin supplementation was chosen based on the identification of essential B vitamins (riboflavin, folic acid and biotin) for *Ornithodoros moubata* (Duron *et al*. 2018).

Based on previous studies, we decided to use a protocol with an antibiotic treatment in the blood meal instead of using antibiotic injections or hosts treated with antibiotics (Duron *et al*. 2018; Guizzo *et al*. 2017; Zhang *et al*. 2017; Ninio *et al*. 2015). While injection is extremely efficient to ensure that the right amount of antibiotics is administered, it remains traumatic for the tick and is not adapted for soft tick species (S. Bonnet & Liu 2012). Besides, as a lab species since decades, *O. moubata* presented excellent results with artificial feeding and this method was chosen for easier use and more accurate control of antibiotic concentrations.

In the first experiment, a total of 60 first stage nymphs were engorged for each treatment condition. A total of 20 individuals were sampled for molecular analyses at day 0 (before blood meal), and 10 individuals at day 1, day 3, day 5, day 10, day 15 and day 30 after blood meal (**Figure 1**). Nymphs which died before DNA extraction were excluded. For more details about both the sample size and group size, please see supplementary data (**Table S1**).

In the second experiment, adult stages (males and females) were engorged with two consecutive treated blood meals with an interval of 52 days between the two. Treatment groups were the same as the one used for nymphs (control, rifampicin, gentamycin, with or without B vitamins).

Each female tick was weighted before and after the blood meal. To ensure efficient blood meal, ticks were fed in groups: four to seven females were fed together with 3 males in one well containing 9 mL of blood. Therefore, we had to mark female ticks individually with different colors using Posca felt pens. If female ticks gained less than 50 mg at the end of the first blood meal, the blood feeding was considered as a failure and ticks were fed again during a second blood meal and a third one if needed. If after three attempts some female ticks were still not fed, they were excluded from the study. Males were only used for mating and were not monitored afterward.

Feeding success and tick weight after blood meal were measured to estimate bias in the success of blood feeding between treated and control groups. No statistical differences were revealed for these two parameters in relation with the antibiotic treatment or vitamin supplementation.

At the end of the experiment (after two treatments and reproduction monitoring), female ticks which were still alive were stored at -80°C for DNA extraction and qPCR analyses.

### DNA extraction

Nymphs and adults were washed 30 seconds in 1% bleach solution and then rinsed for 1 minute in three successive baths of Milli-Q water to eliminate cuticular bacteria (Binetruy *et al*. 2019). Ticks were then crushed individually using a Qiagen TissueLyser instrument with steel and plastic beads in 300 μL of DMEM medium complemented at 10% with calf serum. DNA was extracted from 75 μL of each crushed tick homogenate, using the genomic DNA extraction kit NucleoSpin^®^ Tissue (Macherey-Nagel, Hoerdt, Germany), and following the standard protocol for human or animal tissue and cultured cells. DNA extracts were finally eluted in 50 μl of elution buffer and stored at -20°C until further use.

### Quantitative PCR

Quantitative PCRs were launched using Roche Lightcycler^®^ 96 instruments (Roche Applied Sciences, Beijing, China) using the taqMan method.

Extracted DNA (2 μL) was added to the amplification mix (18 μL: Mix 10 μL, H2O 6.6 μL, forward Primer Actin 0.6 μM, reverse Primer Actin 0.6 μM, Probe Actin 0.2 μM, forward Primer target gene 0.6 μM, reverse Primer target gene 0.6 μM, Probe target gene 0.2 μM). Primers used to amplify *Francisella* targeted the *RpoB* gene coding for the β subunit of RNA polymerase. Primer sequences were Forward: 5’-GTGGTGTACCTATTGCTACG-3’, Reverse: 5’-TGATGGTCGTACTGGTAAGC-3’ and Probe: 5’-ATAGGTGGCTTTGCAACTAA-3’. Primers used to amplify *Rickettsia* targeted the *GltA* gene coding for the citrate synthase enzyme. Primer sequences were Forward: 5’-AATGATCCGTTTAGGTTGATG-3’, Reverse: 5’-ACCCACTCCTGCAAATAG-3’ and Probe: 5’-GTGCCGCAGTACTTAAAGA-3’. The actin gene from *Ornithodoros moubata* was also amplified as a control for the assessment of DNA extraction success and to standardize the results with the tick size. Primer sequences were Forward: 5’-CCGGTATTGCCGACCGTATGC-3’, Reverse: 5’-CTCCCTGTCCACCTTCCAGC-3’ and Probe: 5’-CGAGAGGAAGTACTCCGTCTGG-3’ (Pereira De Oliveira, Vial & Le Potier 2022; Duron *et al*. 2018).

The PCR cycling program was 1 min at 95°C, then 45 cycles of 1 min at 95°C, 30 sec at 56°C. Analyses were performed using both FAM and HEX fluorescence indicators. Samples with a cycle threshold (Ct) beyond 38 were considered as negative. Cts were converted into concentrations from the standard curves performed with dilutions of a DNA plasmid containing the targeted sequences. Results are presented as ratio of the concentration of the gene of interest (GltA or RpoB) divided by the concentration of the actin gene to standardize for the tick size.

Positive and negative controls were included in each qPCR, using 2 μL of DNA of tick previously tested positive for *FLE* and *Rickettsia* (Duron *et al*. 2018) and 2 μL of nuclease-free water respectively. In addition, qPCRs were performed on blood samples (used for blood meal) and on milli-Q water (used for antibiotics and vitamins dilution). They showed no amplification for FLE and Rickettsia indicating that blood meal and milli-Q water were not contaminated with those bacteria.

### Reproduction monitoring

After treated blood meal, females were kept in the same tubes as males for seven days to allow mating, then each female was isolated in an individual plastic tube. The monitoring of the tick reproduction was blinded regarding applied treatments. A total of 132 females were included in the study and monitored every day during two successive 50-days periods after blood meal. For each tick, the laying date, the number of eggs laid and the egg mass were recorded. The number of eggs laid was standardized with the weight of the female tick after blood meal. For each egg, the date of hatching and of molting into first nymphal stage were recorded (**Figure 1**).

### Statistical analysis

Statistical analyses were performed using R version 4.2.3 (2023-03-15) to test the effect of antibiotic and vitamin treatments on DNA concentrations and traits associated with reproductive fitness (R Core Team 2023). For the analyses of qPCR results, ratios of DNA concentrations were log-transformed and generalized least-squares (GLS) models were used with the function **gls** to allow different variances for each level of the explanatory variables (Zuur *et al*. 2009). The best model including optimal variance structure and fixed effects (namely, antibiotic and vitamin treatments and their interaction) was determined by AIC comparisons. In nymphs, due to material constraints, two different experimenters performed the qPCR at two different moments. This was added as a fixed effect in all the models and had a clear effect on measured ratios (GltA: p=1.60×10-05, F-value=19.2 and, RpoB: p=1.14×10-65, F-value=473). DNA concentrations were also log-transformed and analyzed using **glm** with a gamma distribution when ratios were not relevant for the analyzes. To analyze reproduction parameters, we used generalized linear models with the function **lm, glm** or generalized mixed model with **glmer** (R package **Lme4** (Bates *et al*. 2015)) according to the distribution of the data and the absence (**glm**) or presence (**glmer**) of a random effect. The random effect was added for analyses targeting the offspring of tick females to account for the lack of independence of data points (since measured individuals may be offspring of the same mother). For egg incubation time and time for molting into first nymphal stage, the level of variability between mothers was not sufficient to warrant incorporating it as a random effect. Depending on the variable, normal, Poisson or binomial distributions were used (see **Table 1**). Overdispersion and underdispersion of the data were estimated using a quasi-Poisson or quasi-binomial distribution. Statistical comparisons were performed using either F-test, chi-squared test or Kruskal-Wallis test. Post-hoc analyses were conducted using the function **emmeans** (Tukey HSD test) to identify statistically distinct treatment groups. When the mean was presented for a trait, standard deviation (SD) was indicated in brackets. In addition, data from the two 50-days periods were statistically compared and analyzed using the models described above. Interactions with antibiotic treatments and vitamin B supplementation were taken into account in the analysis.

**Table 1:** Statistical analyses performed on traits associated with reproductive fitness measured in female ticks during 50 days after the first antibiotic treatment and vitamin supplementation.

| Variable | Family | Effect of the treatment | Effect of the vitamin | Effect of Interaction |
| --- | --- | --- | --- | --- |
| Survival over 50 days after blood meal | Binomial | Pr: 0.00969<br>Dev: 9.27 | Pr: 0.377<br>Dev: 0.781 | Pr: 0.187<br>Dev: 3.36 |
| Laying probability | Binomial | Pr: 0.350<br>Dev: 2.10 | Pr: 0.107<br>Dev: 2.59 | Pr: 0.448<br>Dev: 1.61 |
| Number of days to first oviposition | Quasi-Poisson<sup>1</sup> | Pr: 0.374<br>Dev: 3.55 | Pr: 0.0582<br>Dev: 6.47 | Pr: 0.161<br>Dev: 6.59 |
| Number of eggs standardized by mother weight<sup>3</sup> | Quasi-Poisson<sup>1</sup> | Pr: 0.412<br>Dev: 18.7 | Pr: 0.770<br>Dev: 0.902 | Pr: 0.917<br>Dev: 1.82 |
| Average egg mass | Normal | Pr: 0.353<br>Dev: 0.00227 | Pr: 0.144<br>Dev: 0.00468 | Pr: 0.889<br>Dev: 0.000253 |
| Egg incubation time | Quasi-Poisson<sup>2</sup> | Pr: 0.696<br>Dev: 0.0617 | Pr: 0.765<br>Dev: 0.00759 | Pr: 0.0570<br>Dev: 0.488 |
| Egg hatching rate | Binomial | Pr: 0.748<br>Dev: 0.582 | Pr: 0.957<br>Dev: 0.00295 | Pr: 0.550<br>Dev: 1.20 |
| Time for molting into first nymphal stage | Quasi-Poisson<sup>2</sup> | Pr: 0.710<br>Dev: 0.0470 | Pr: 0.392<br>Dev: 0.0501 | Pr: 0.744<br>Dev: 0.0405 |
| Molting rate | Binomial | Pr: 0.871<br>Dev: 0.276 | Pr: 0.184<br>Dev: 1.76 | Pr: 0.763<br>Dev: 0.542 |

Note: Pr: P-value. Dev: Deviance, $\chi^2$ for binomial and quasi-Poisson distribution, F-value for normal distribution. Degree of freedom was 2 for the effect of the treatment and the effect of the interaction and 1 for the effect of vitamin. <sup>1</sup> Distribution corrected for overdispersion of the data. <sup>2</sup> Distribution corrected for underdispersion of the data. <sup>3</sup> Variable corrected using an offset of the tick weight. Statistically significant results are highlighted with a grey background.

### Ethics

The use of cows which provided blood for tick engorgement complied with the ethical standards of European regulations (directive from the European Council from November 24^th^, 1986 (86/609 / EEC)) and French legislation (ethic committee CEEA-LR-36 available on the APAFIS platform) on the care and use of animals. More precisely, blood sampling has been recorded in 2021 under the agreement number APAFIS#27628-2020101010475151 for the CIRAD-Montpellier rearing facility G 34-172-15.

## Results

### Molecular results

A total of 366 first stage nymphs were tested for *Rickettsia* (target gene *GltA*) and *Francisella*-like endosymbiont (target gene *RpoB*).

Out of 366 samples, 98.6% were tested positive for *Rickettsia* (359/364) while two samples were excluded due to the abnormal qPCR curves. Two nymphs were tested negative in the control group without vitamins (at day 3 and day 5) and three others in the gentamycin group (day 30 with and without vitamins and day 10 without vitamins). The concentration of *GltA* DNA increased over time after blood meal in the control (p=0.00353, 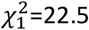) and gentamycin (p=4.74×10-11, 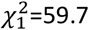) groups but not in the rifampicin group (p=0.262, 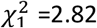). Similar increase also appeared for the actin gene in all groups (p<2.20×10-16, 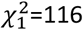).

After standardization, the ratio of DNA concentration remained significantly lower in the rifampicin group than in the two other groups (p=1.62×10-92, F-value=429), independently from vitamin supplementation (**Figure 2A and 2B**).

**Figure 2:**
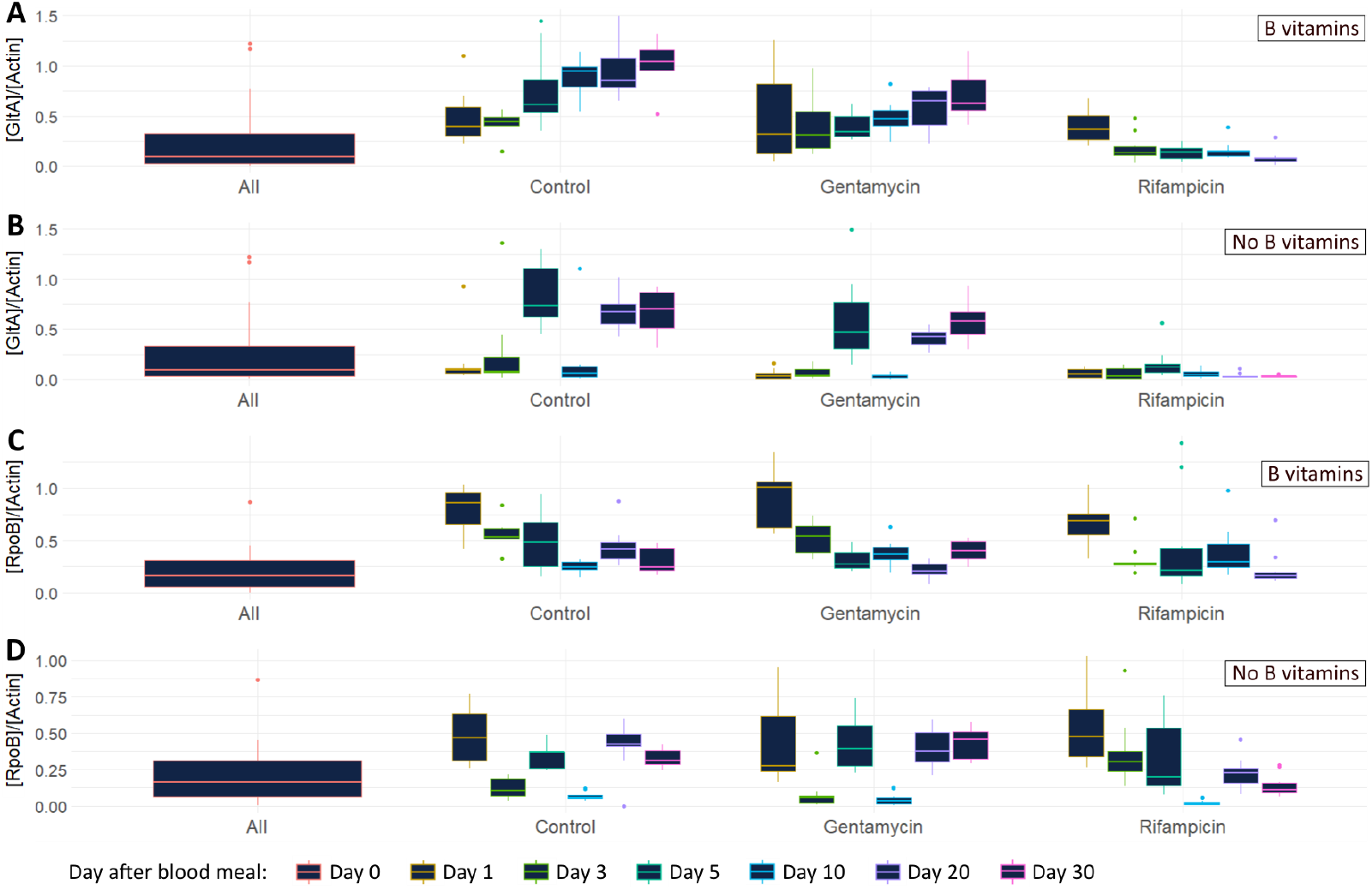
Ratios of DNA concentration obtained by qPCR for *Rickettsia* and *Francisella*-like endosymbiont (FLE) after antibiotic treatment of individual first stage nymphs. **(A)** Ratio of the concentration of *GltA* DNA ([GltA]), indicator of the presence of *Rickettsia* over the concentration of *O. moubata* actin DNA [Actin]. Nymphs supplemented with B vitamins. **(B)** Ratio of [GltA] over [Actin]. Nymphs not supplemented with B vitamins. **(C)** Ratio of the concentration of *RpoB* DNA ([RpoB]), indicator of the presence of FLE over [Actin]. Nymphs supplemented with B vitamins **(D)** Ratio of [RpoB] over [Actin]. Nymphs not supplemented with B vitamins.

Regarding *Francisella*-like endosymbiont, 98.1% of the qPCR were positive (359/366) with one negative samples at day 0, two in the rifampicin group (day 1 no vitamin, and day 10 with vitamin) and four in the control group (day 5 and 10 with vitamin, and day 20 and 30 no vitamin). As for *Rickettsia*, there was an increase in the quantity of *RpoB* DNA detected over time in the gentamycin group (p=0.00309, 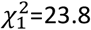) but not in the control (p=0.0869, 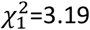) and rifampicin (p=0.134, 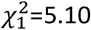) groups. After standardization, the ratio of DNA concentration remained significantly different between the three treatment groups (p=1.71×10-12, F-value=29.4) (**Figure 2C and 2D**).

At the end of the reproduction monitoring, a total of 59 females were still alive and were collected for quantification of *Rickettsia* and *Francisella*-like endosymbionts. Forty-eight females were tested positive for *Rickettsia* (14/14 control, 2/6 gentamycin without vitamin, 11/12 gentamycin with vitamin, 10/14 rifampicin without vitamin, and 11/13 rifampicin with vitamin), all females were tested positive for *Francisella*-like endosymbiont, and five adults were excluded due to abnormal qPCR curves. Comparing the ratio of concentration of symbionts, no significant differences were observed between the groups concerning vitamin supplementation (p>0.05). However, focusing on the different antibiotic treatment groups, it appeared that ticks treated with rifampicin or gentamycin hosted lower concentrations of *Rickettsia* (p=1.81×10-06, F-value=19.1) and *Francisella*-like endosymbiont (p=1.12×10-09, F-value= 31.7) than ticks from the control group. This effect was stronger in the rifampicin group than in the gentamycin group (**Figure 3A and 3B**).

**Figure 3:**
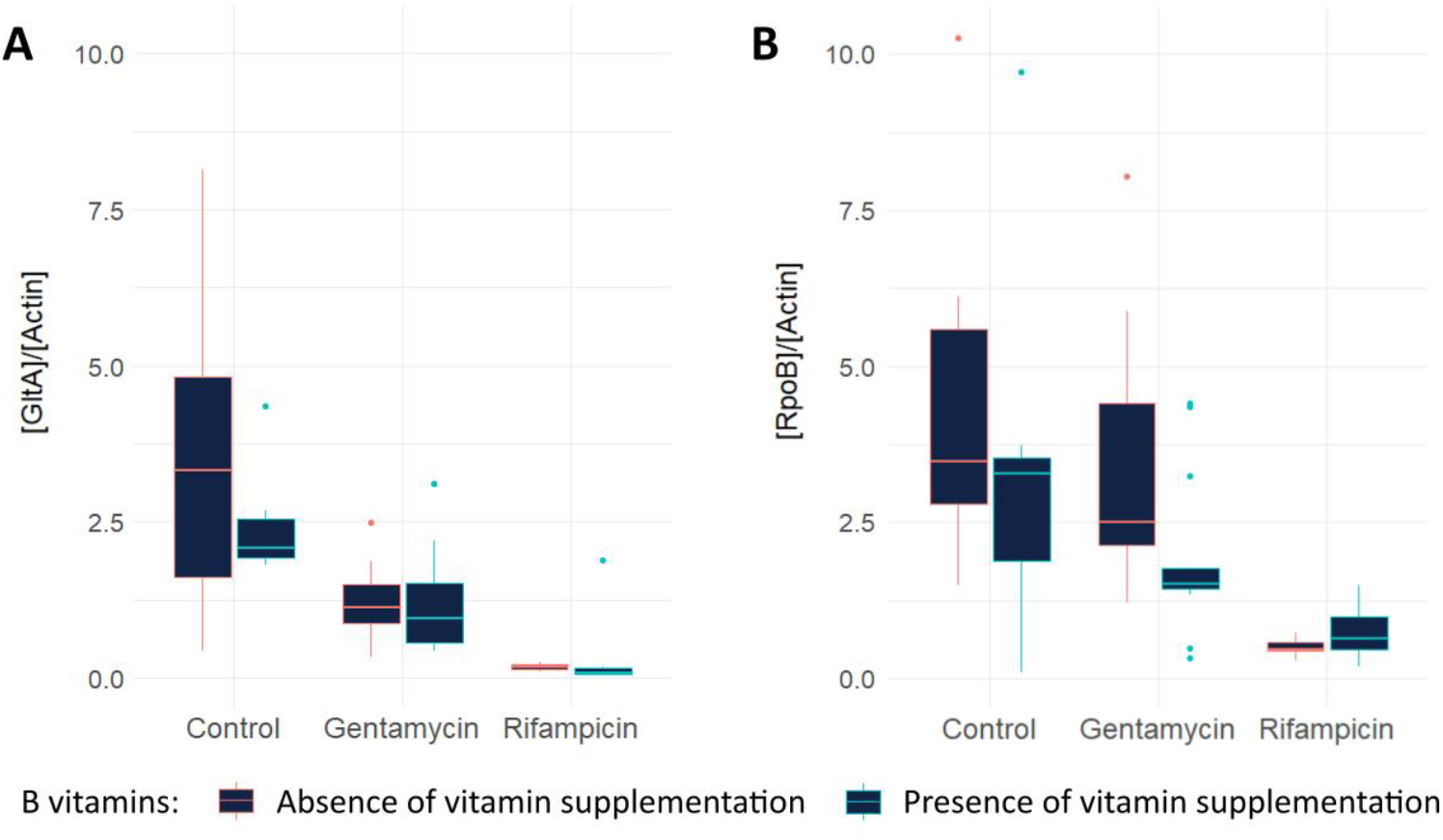
Ratio of DNA concentration obtained by qPCR for *Rickettsia* and *Francisella*-like endosymbiont (FLE) after antibiotic treatments of female ticks. **(A)** Ratio of the concentration of *GltA* DNA ([GltA]), indicator of the presence of *Rickettsia* over the concentration of *O. moubata* actin DNA [Actin]. **(B)** Ratio of the concentration of *RpoB* DNA ([RpoB]), indicator of the presence of FLE over [Actin]. Ticks were sampled after two consecutive treatments and monitoring of the reproduction (50 days after the second treatment).

### Monitoring of the reproduction

Adult ticks were separated in three treatment groups with two subgroups in each (half with vitamin B supplementation and half without). Effects of antibiotic treatment, vitamin supplementation and the interaction between the two were evaluated for statistical differences on traits associated with reproductive fitness over two successive periods of 50 days (**Table 1 and 2**).

**Table 2:** Statistical analyses performed on traits associated with reproductive fitness measured in female ticks during 50 days after the second antibiotic treatment and vitamin supplementation.

| Variable | Family | Effect of the treatment | Effect of the vitamin | Effect of Interaction |
| --- | --- | --- | --- | --- |
| Survival over full duration of the experiment (102 days) | Binomial | Pr: 0.00655<br>Dev: 10.1 | Pr: 0.960<br>Dev: 0.00250 | Pr: 0.0672<br>Dev: 5.40 |
| Survival over 50 days after the second blood meal | Binomial | Pr: 0.150<br>Dev: 3.79 | Pr: 0.711<br>Dev: 0.138 | Pr: 0.0482<br>Dev: 6.07 |
| Laying probability | Binomial | Pr: 0.0890<br>Dev: 5.15 | Pr: 0.729<br>Dev: 0.128 | Pr: 0.247<br>Dev: 2.97 |
| Number of days to first oviposition | Quasi-Poisson<sup>1</sup> | Pr: 0.682<br>Dev: 4.21 | Pr: 0.132<br>Dev: 12.5 | Pr: 0.208<br>Dev: 17.2 |
| Number of eggs standardized by mother weight<sup>3</sup> | Quasi-Poisson<sup>1</sup> | Pr: 0.259<br>Dev: 132 | Pr: 0.304<br>Dev: 51.8 | Pr: 0.0171<br>Dev: 398 |
| Average egg mass | Normal | Pr: 0.987<br>Dev: 0.0000662 | Pr: 0.861<br>Dev: 0.000151 | Pr: 0.360<br>Dev: 0.00512 |
| Egg incubation time | Quasi-Poisson<sup>2</sup> | Pr: 0.325<br>Dev: 0.250 | Pr: 0.680<br>Dev: 0.0189 | Pr: 0.0157<br>Dev: 0.924 |
| Egg hatching rate | Binomial | Pr: 0.736<br>Dev: 0.615 | Pr: 0.190<br>Dev: 1.72 | Pr: 0.480<br>Dev: 1.47 |
| Time for molting into first nymphal stage | Quasi-Poisson<sup>2</sup> | Pr: 0.000383<br>Dev: 1.26 | Pr: 0.625<br>Dev: 0.0192 | Pr: 0.901<br>Dev: 0.0168 |
| Molting rate | Binomial | Pr: 0.210<br>Dev: 3.12 | Pr: 0.717<br>Dev: 0.131 | Pr: 1.00<br>Dev: 6.72×10<sup>-5</sup> |

Note: Pr: P-value. Dev: Deviance, X<sup>2</sup> for binomial and quasi-Poisson distribution, F-value for normal distribution. Degree of freedom was 2 for the effect of the treatment and the effect of the interaction and 1 for the effect of vitamin. <sup>1</sup> Distribution corrected for overdispersion of the data. <sup>2</sup> Distribution corrected for underdispersion of the data. <sup>3</sup> Variable corrected using an offset of the tick weight. Statistically significant results are highlighted with a grey background.

After the first treatment, there was no effect of antibiotic treatment on traits associated with reproductive fitness, except for survival (Figure 4A). Survival was significantly different between the three treatment groups but not between the vitamin subgroups (**Figure 4A, Table 1**), with the control group presenting a lower survival rate (65.2%) than the gentamycin (90.9%) group (p=0.0165). Throughout the first period of monitoring, 78.8% (104/132) of the female ticks survived.

**Figure 4:**
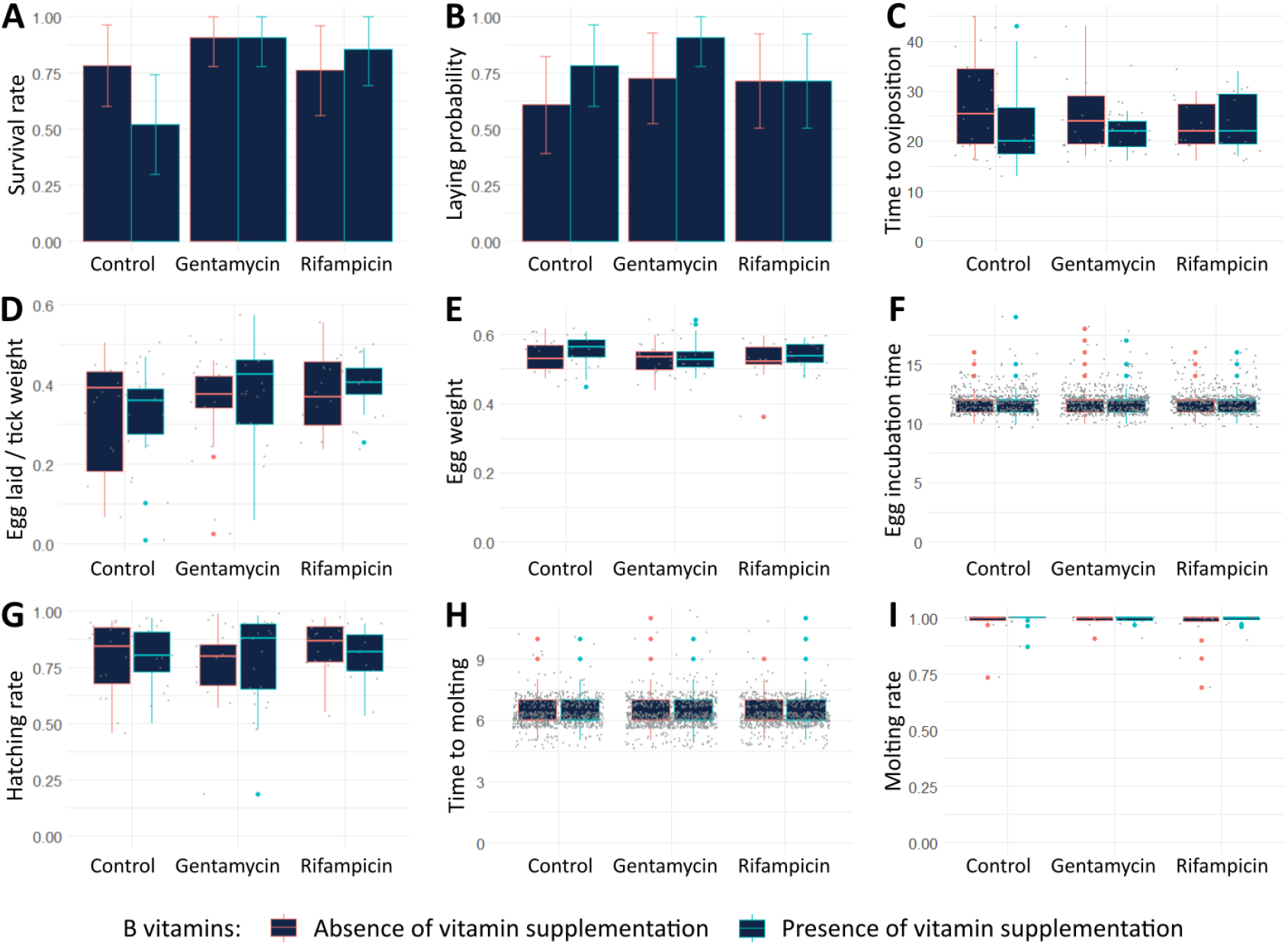
Traits associated with reproductive fitness measured in female ticks during 50 days after the first antibiotic treatment and vitamin supplementation. **(A)** Tick survival rate. **(B)** Laying probability: number of ticks laying eggs/ population size. **(C)** Time between blood meal and first day of oviposition (in days). **(D)** Number of eggs laid per tick corrected for the mass of the blood fed female tick (number of eggs per mg of tick weight). **(E)** Mean egg weight (in mg). **(F)** Egg incubation time (in days). **(G)** Egg hatching rate. **(H)** Time between hatching and molting into first nymphal stage (in days) **(I)** Molting success rate. In histograms, 95% confidence intervals are presented.

During these first 50 days, 74.2% (98/132) of engorged ticks initiated oviposition and the mean duration between blood meal and the onset of oviposition was 23.9 days [SD: 6.81] (**Figure 4B, Figure 4C**). These ticks laid between 2 and 220 eggs, during the first 50 days of monitoring, with an average of 109 eggs (or, after standardization, 0.361 egg per mg of tick weight [SD: 0.116]) and an egg mass of 0.538 mg [SD: 0.0463] (**Figure 4D, Figure 4E**).

Mean values for developmental parameters were 79.6% of hatching success [SD: 15.1] with an egg incubation time of 11.9 days [SD: 1.01] (**Figure 4F, Figure 4G**) and 98.4% for molting rates [SD: 4.77] with 6.30 days to molt into first nymphal stage [SD: 0.657] (**Figure 4H, Figure 4I**).

After the first period of 50 days, the surviving ticks were fed a second time with the same feeding protocol. Female ticks were monitored during a new period of 50 days and parameters associated with reproduction were measured (**Table 2**).

Measurements after the second blood meal yielded largely similar results (**Figure 5, Table 2, Table S2**). There was no effect of antibiotic and vitamin treatments, except for the time before molting which was increased for larvae of the rifampicin group (6.59 days against 6.38 for the control group and 6.34 for the gentamycin group, p=0.00471). No effect was observed regarding vitamin supplementation (**Figure 5H, Table 2**).

**Figure 5:**
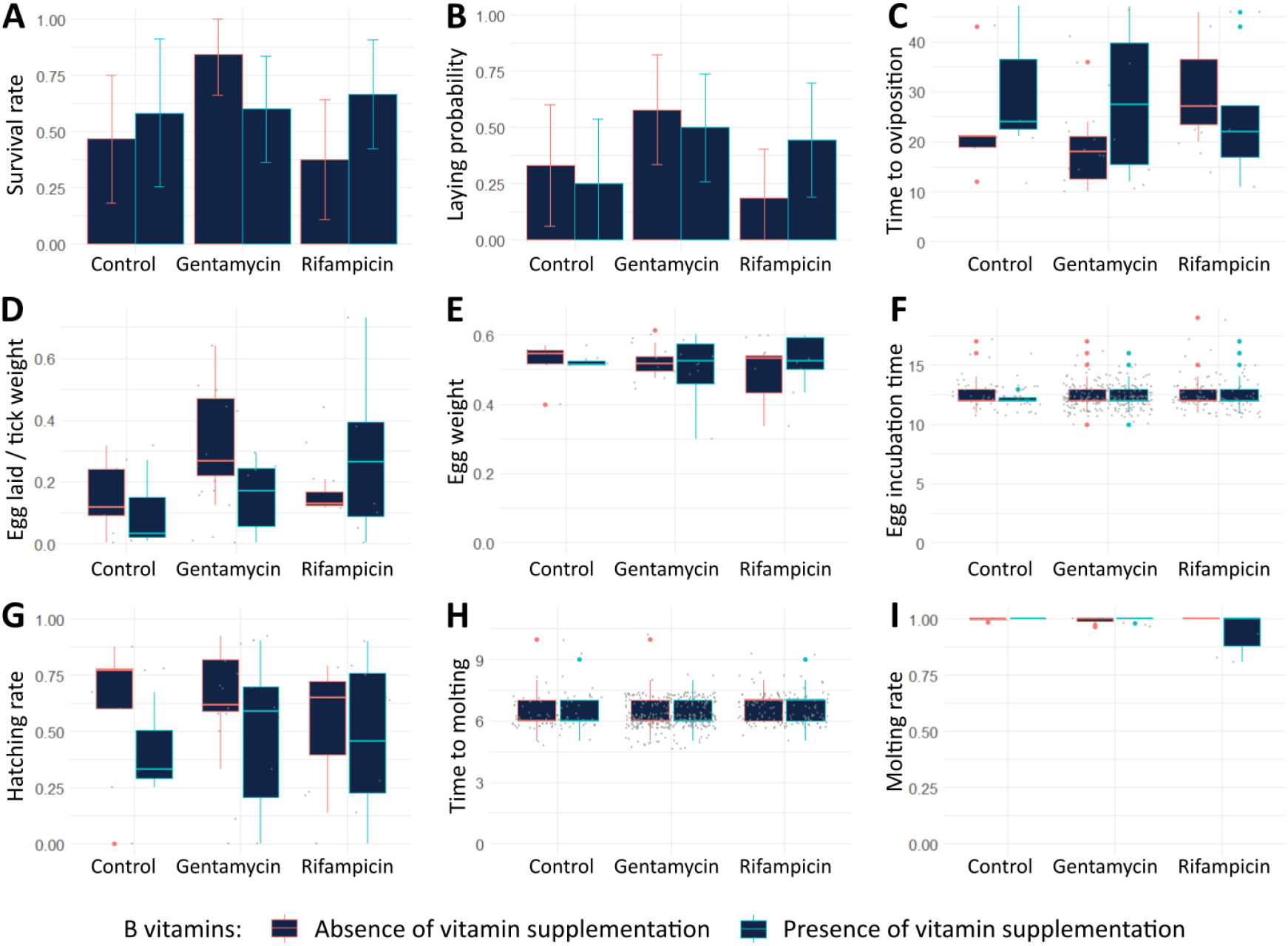
Traits associated with reproductive fitness measured in female ticks during 50 days after the second antibiotic treatment and vitamin supplementation. **(A)** Tick survival rate. **(B)** Laying probability: number of ticks laying eggs/ population size. **(C)** Time between blood meal and first day of oviposition (in days). **(D)** Number of eggs laid per tick corrected for the mass of the blood fed female adult tick (number of eggs per mg of tick weight). **(E)** Mean egg weight (in mg). **(F)** Egg incubation time (in days). **(G)** Egg hatching rate. **(H)** Time between hatching and molting into first nymphal stage (in days) **(I)** Molting success rate. In histograms, 95% confidence intervals are presented.

Considering the whole experiment (two blood meals), a significant effect of the antibiotic treatment group on survival is observed (control group versus gentamycin group, p=0.00920). The control groups presented some of the lowest survival rates (33.3% and 30.4% respectively in absence and presence of vitamins). The rifampicin group without B vitamins also presented a low survival rate (30.0%), whereas the three other groups presented higher survival rates: 57.1% for rifampicin with vitamin, 54.5% and 76.2% for the gentamycin group with or without vitamins (**Figure 6, Table 2)**.

**Figure 6:**
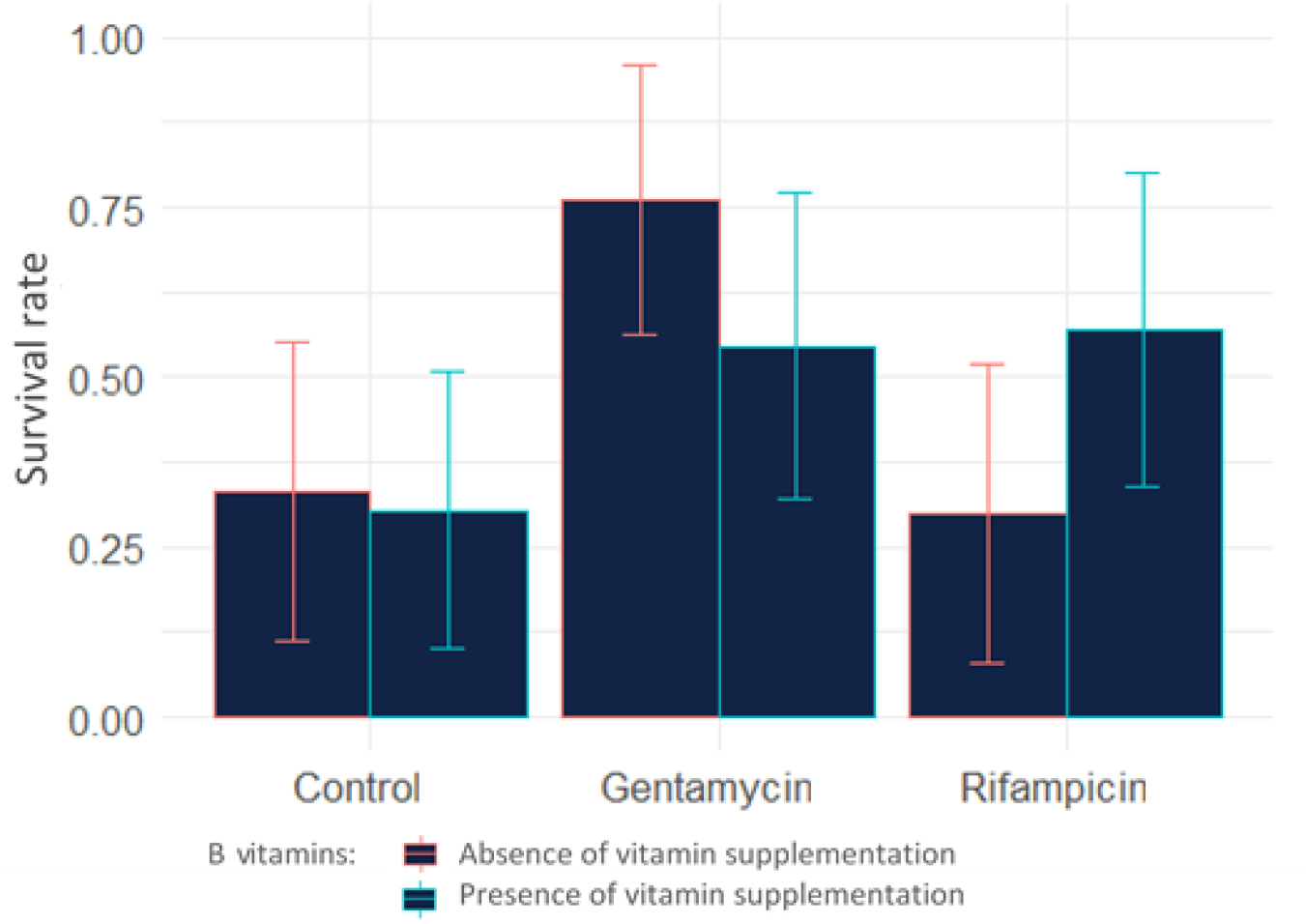
Survival rate of *Ornithodoros moubata* female ticks after antibiotic treatments and vitamin supplementation over the whole experiment (102 days). 95% confidence intervals are presented.

Following the ticks over two consecutive blood meals brings information on how the tick reproduction parameters are affected over time in *Ornithodoros moubata* (**Table 3, Table S2**). Almost all traits associated with reproductive fitness were negatively affected during the second 50-days period after blood meal: survival rates, laying probability, number of eggs laid, egg mass and hatching rate decreased whereas egg incubation time and molting time increased. The number of days to oviposition and the molting rate did not change after the second blood meal. These observations are mostly independent from the treatment received during blood meals.

**Table 3:**
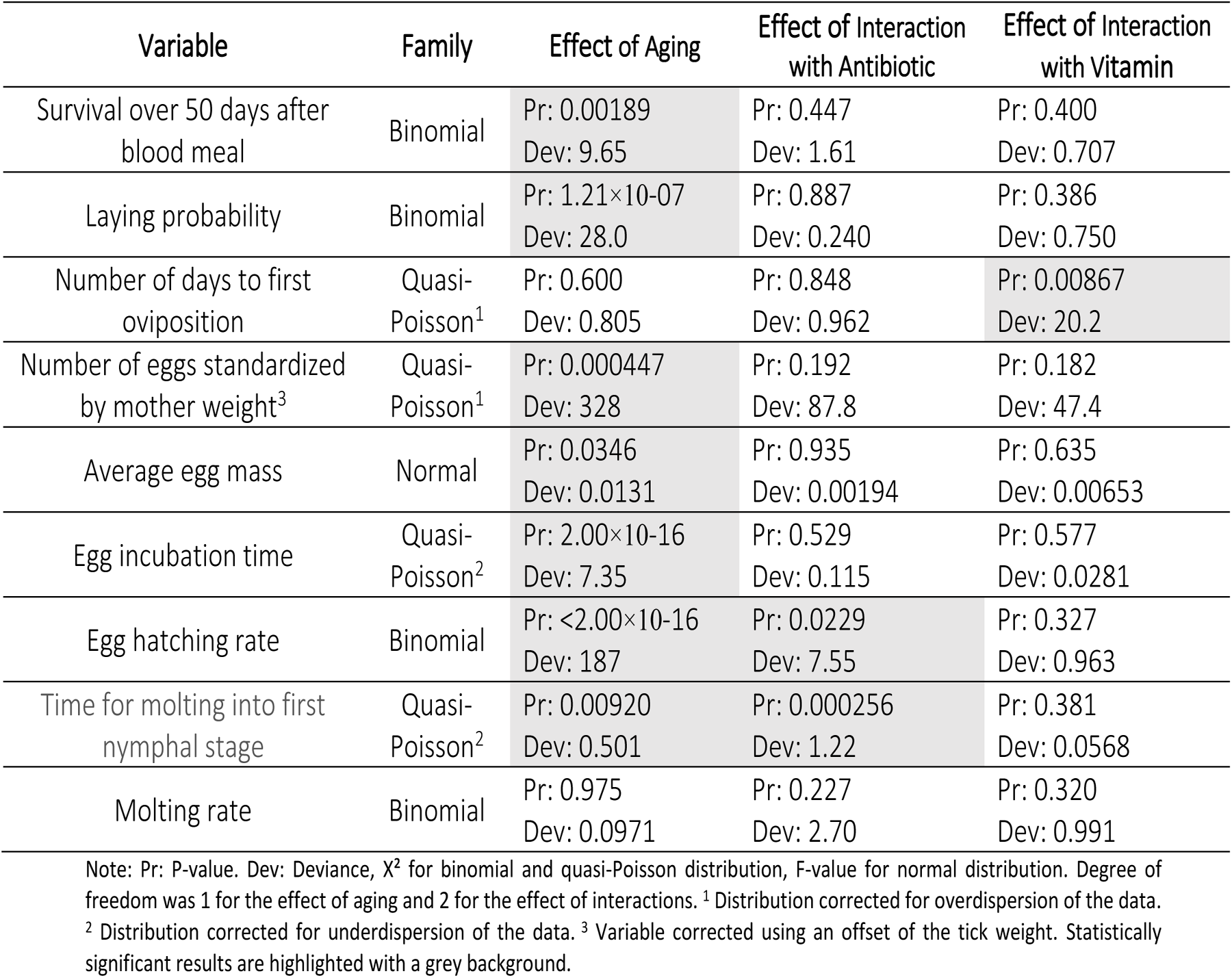
Comparison of the reproductive indicators measured in adult female ticks between the two 50 days periods after blood meal with antibiotic treatments (gentamycin, rifampicin, or negative control) and vitamin supplementation (presence of vitamins and absence of vitamins).

| Variable | Family | Effect of Aging | Effect of Interaction with Antibiotic | Effect of Interaction with Vitamin |
| --- | --- | --- | --- | --- |
| Survival over 50 days after blood meal | Binomial | Pr: 0.00189<br>Dev: 9.65 | Pr: 0.447<br>Dev: 1.61 | Pr: 0.400<br>Dev: 0.707 |
| Laying probability | Binomial | Pr: 1.21×10-07<br>Dev: 28.0 | Pr: 0.887<br>Dev: 0.240 | Pr: 0.386<br>Dev: 0.750 |
| Number of days to first oviposition | Quasi-Poisson<sup>1</sup> | Pr: 0.600<br>Dev: 0.805 | Pr: 0.848<br>Dev: 0.962 | Pr: 0.00867<br>Dev: 20.2 |
| Number of eggs standardized by mother weight<sup>3</sup> | Quasi-Poisson<sup>1</sup> | Pr: 0.000447<br>Dev: 328 | Pr: 0.192<br>Dev: 87.8 | Pr: 0.182<br>Dev: 47.4 |
| Average egg mass | Normal | Pr: 0.0346<br>Dev: 0.0131 | Pr: 0.935<br>Dev: 0.00194 | Pr: 0.635<br>Dev: 0.00653 |
| Egg incubation time | Quasi-Poisson<sup>2</sup> | Pr: 2.00×10-16<br>Dev: 7.35 | Pr: 0.529<br>Dev: 0.115 | Pr: 0.577<br>Dev: 0.0281 |
| Egg hatching rate | Binomial | Pr: <2.00×10-16<br>Dev: 187 | Pr: 0.0229<br>Dev: 7.55 | Pr: 0.327<br>Dev: 0.963 |
| Time for molting into first nymphal stage | Quasi-Poisson<sup>2</sup> | Pr: 0.00920<br>Dev: 0.501 | Pr: 0.000256<br>Dev: 1.22 | Pr: 0.381<br>Dev: 0.0568 |
| Molting rate | Binomial | Pr: 0.975<br>Dev: 0.0971 | Pr: 0.227<br>Dev: 2.70 | Pr: 0.320<br>Dev: 0.991 |

Note: Pr: P-value. Dev: Deviance, X<sup>2</sup> for binomial and quasi-Poisson distribution, F-value for normal distribution. Degree of freedom was 1 for the effect of aging and 2 for the effect of interactions. <sup>1</sup> Distribution corrected for overdispersion of the data. <sup>2</sup> Distribution corrected for underdispersion of the data. <sup>3</sup> Variable corrected using an offset of the tick weight. Statistically significant results are highlighted with a grey background.

## Discussion

In this study, we used antibiotic treatments to eliminate key members of the microbiota of the tick *O. moubata* and evaluate the consequences of this dysbiosis on several parameters of the tick biology. We thus focused our study on the two main endosymbionts of *O. moubata* known to be crucial for the tick biology: *Francisella-*like endosymbionts (FLE) and *Rickettsia*.

### Impacts of antibiotics on the tick microbiota

The qPCR results suggest that the rifampicin treatments limited the growth and/or decreased the population size for *Rickettsia* and FLE. It seemed that the effect was bacteriostatic after one treatment on nymphs during growth. The diminution of the population of symbionts after two treatments in adult females can be the consequence of this bacteriostatic effect over time or the outcome of a bactericide effect after several consecutive treatments. In the end, the treatment performed was sufficient to prevent the growth of FLE and *Rickettsia* even though insufficient to eliminate them completely.

The gentamycin treatment induced only a slight decrease on the ratio of DNA concentration related with FLE. It seems that this antibiotic or, at least, the posology used for this antibiotic, was less effective than the rifampicin. Besides, the lack of effect of the gentamycin on *Rickettsia* populations was expected since *Rickettsia* are known to be resistant to this antibiotic (Vanrompay *et al*. 2018). Finally, no effect of the vitamin group was reported on the concentration of endosymbiont in adult ticks. This result was expected as endosymbionts do not depend on B vitamins, they rather produce B vitamins.

Moreover, in this experiment, a significant increase in the amount of bacterial DNA, for both *Francisella*-like endosymbionts and *Rickettsia*, was observed during 30 days after blood meal. It also appeared that the amount of actin DNA increased over the same period of time, suggesting a strong metabolic activity with multiple cellular divisions. This is most probably linked to the molting process which takes 10.5 days [SD: 1.06] under those conditions (Jourdan-Pineau, unpublished data). As intracellular bacteria, it is likely that *Francisella* and *Rickettsia* took advantage of this metabolic increase for their own growth as observed in other species (Sassera *et al*. 2008). These endosymbionts might be simply more active during this period of intense metabolic activity which would be in agreement with the alterations of the molting process observed after antibiotic treatments (Duron *et al*. 2018).

### Consequences of the modification of the microbiota on the tick physio-biology

In female ticks, considering only the antibiotic treatments, no effect was detected on the reproduction itself after the first treatment. This result would suggest that the FLE and *Rickettsia* are not involved in the tick reproduction. However, at this point of the experiment it is also possible that the effect of antibiotic treatments did not eliminate enough symbiont yet. In parallel, we observed that the negative control group presented a higher mortality rate than the antibiotic groups. This surprising result led us to consider the fitness cost of harboring endosymbionts, as already referenced for other arthropods (Ankrah, Chouaia & Douglas 2018). For instance, it is possible to hypothesize that *Francisella* provides necessary nutrients for ticks and helps them avoid deficiencies in the long term; however, this may come at a short-term cost, resulting in increased tick mortality during reproduction.. Another possible explanation would be that other bacteria than FLE and *Rickettsia* were eliminated by the antibiotic treatments and that those bacteria presented a fitness cost for the tick. This hypothesis remains less probable than a role of FLE and Rickettsia which are the most frequent bacteria with key roles in the development and survival of *O. moubata* (Duron *et al*. 2018).

When looking at vitamin supplementation, no effect was detected on the reproduction or survival after the first treatment. It is possible that the duration of our experiment was not sufficient to fully observe the effects of a vitamin deficiency. Moreover, FLE were maintained at a low concentration but not completely eliminated here, suggesting that all ticks were already supplied, at some extent, in B vitamins.

If we consider the results obtained after the second antibiotic treatment, two main elements can be stressed out. First, is the confirmation of an effect on tick survival. After both treatments, the control group presented a higher mortality than the gentamycin group. However, after the second treatment, the rifampicin group was also affected by an increased mortality rate. One hypothesis could be that more FLE have been eliminated after the second treatment, resulting in a more severe vitamin B deficiency in ticks compared to the first treatment, which in turn caused the death of adult female ticks. This hypothesis is comforted by the fact that the survival rate in the rifampicin group after the second treatment was better when the ticks were supplemented with B vitamins. From these results, it would seem that there is a cost to maintain high levels of symbionts (increased mortality in the control group), but the absence of symbionts results in a similar death rate (mortality in the second rifampicin group), and finally in the groups with moderate amount of symbiont, the benefits of endosymbiont compensate any associated fitness cost (gentamycin treatment groups).

Second, when looking at the parameters associated with reproduction, no clear-cut effect can be established after the second treatment. The few significant results (time to molting) presented variations of low range. The variations on the molting time are oriented with a longer time period for the rifampicin group. As suggested in the literature, endosymbionts are crucial during molting (Duron *et al*. 2018), and their elimination through rifampicin could indeed lead to a deterioration of the parameters associated with molting. When looking at past literature on the role of endosymbionts in tick reproduction, several effects were associated with the elimination of *Coxiella*-like endosymbionts (CLE) in hard ticks, including a decrease in number of egg laid (Guizzo *et al*. 2017; Zhang *et al*. 2017), an increase in the egg incubation time (Li, Zhang & Zhu 2018), a decrease in the hatching rate (Guizzo *et al*. 2017; Li, Zhang & Zhu 2018; Zhong, Jasinskas & Barbour 2007) and a decrease in the molting success (Zhong, Jasinskas & Barbour 2007). Interestingly, no effects were reported for *Rickettsia* (Kurlovs *et al*. 2014) or FLE. This is especially interesting as all these three endosymbionts have a nutritional role. The results presented here indicate no effect of FLE or *Rickettsia* on the tick reproduction. It is possible that CLE are the only tick symbionts which improve the reproductive fitness of their host in addition to vitamin B production, leading them to remain one of the most commonly found endosymbiont in ticks (Duron *et al*. 2017).

In our study, we presented results issued from two successive treatments. Interestingly, in *O. moubata*, the vitellogenesis processes start during the last nymphal stages (Sonenshine 1991) suggesting that the role of the microbiota on reproduction may occur before adult stage. In the future, antibiotic treatments will have to be performed starting from late nymph stages to evaluate the role of the microbiota during these stages on the reproduction of the adults obtained.

However, blood meal and mating are also critical steps for vitellogenesis processes (Horigane *et al*. 2010) and most studies performed on tick reproduction targeted directly adult females (Guizzo *et al*. 2017; Li, Zhang & Zhu 2018; Zhang *et al*. 2017; Zhong, Jasinskas & Barbour 2007; Kurlovs *et al*. 2014). Parameters such as the number of eggs laid depend directly on the efficiency of the blood meal. Moreover, as demonstrated here and in previous studies (Sassera *et al*. 2008), endosymbiont load increases after the blood meal, supporting the hypothesis that they may have a role during and/or after the meal. Ideally, using germ-free tick lineages during blood meal would provide significant insight into this issue. Such lineages are however difficult to sustain, and antibiotic treatment remains, for now, the main tool available.

Finally, even though it was not the objective of the study, monitoring adult females over two periods of egg-laying brought data on the effects of ageing on the reproduction. Similarly to results obtained in model invertebrates (Tatar 2010), the reproductive indicators in soft ticks were strongly deteriorated between the first and second period of egg-laying. Our results indicate no link between this phenomenon and the presence or absence of FLE and *Rickettsia* suggesting the effect is solely due to ageing.

### Implications for future studies

Our experiments question the use of antibiotics during artificial blood feeding of ticks. In hard ticks, antibiotics and antifungal molecules are often added to the blood to prevent microbial development during the long periods of time needed for the blood meal (Krull *et al*. 2017; Militzer *et al*. 2021; González, Bickerton & Toledo 2021). This use of antibiotics and the molecules chosen should be carefully considered in experimental design as they will modify the tick microbiota and, potentially, affect the tick’s ability to develop normally and to be maintained in a colony. This attention should also be increased for lab experiments designed to identify the potential influence of members of the microbiota on the presence of pathogens and *vice versa*.

Moreover, although we focused on two primary endosymbionts, antibiotic treatments likely reshaped the entire tick microbiota. Eliminating FLE and *Rickettsia* proved challenging, suggesting that endosymbionts are not easily lost. However commensal bacteria from the microbiota may be more susceptible to the treatment. Even though not essential for the tick, these commensal bacteria could harbor secondary functions worth studying, especially regarding the tick’s vector competence to pathogens.

Finally, in this study, offspring of treated mothers showed few signs of impaired or delayed development from larvae to first stage nymphs. However, with these results, we cannot rule out that the microbiota of the offspring might be modified due to the treatment of their mothers. Further experiments on those offspring would allow to characterize their microbiota and observe if they suffer any developmental deficiency during subsequent nymphal stages. A way to test this hypothesis would be to monitor over several consecutive blood meals the offspring of both antibiotic treated and control females.

To conclude, this study suggests that FLE and *Rickettsia* endosymbionts have no direct effect on the reproduction of their tick host. Several trends associated with these bacterial species were highlighted including a potential fitness cost, and strong load increase following blood meal.

However, elucidating the role of the tick microbiota remains an arduous task. Antibiotic treatments constitute a powerful tool for this aim, but protocols need to be adjusted and modified in each tick lineage to take into account the microbiota composition and sensitivity.

## Author contributions

FT performed the experiments described, analyzed the data, and wrote the manuscript. TP provided the material, obtained the funding, and conceived and coordinated the study. MD provided the material and helped perform the experiments. LG helped perform the experiments and analyze the data. HJ provided the material, obtained the funding, conceived and coordinated the study, performed the experiments and analyzed the data. All authors read and corrected the final manuscript.

## Acknowledgements

The authors would like to thank Olivier Duron, Rémi Pereira, and the members of the UMR ASTRE for the discussions on molecular biology protocols, Facundo Muñoz for the discussions about statistics as well as Mélanie Jeanneau for her help on DNA extractions.

All experiments on arthropods have been performed on the Baillarguet insectarium platform (https://doi.org/10.18167/infrastructure/00001),member of the National Infrastructure EMERG’IN and of the Vectopole Sud network (http://www.vectopole-sud.fr/). The Baillarguet insectarium platform is led by the joint units Intertryp (IRD, Cirad) and ASTRE (Cirad, INRAE).

## Data, scripts and codes availability

Data are available online: https://doi.org/10.5061/dryad.bg79cnpfm.

Scripts are available online: https://doi.org/10.5281/zenodo.7937778.

## Supplementary material

**Table S1:**
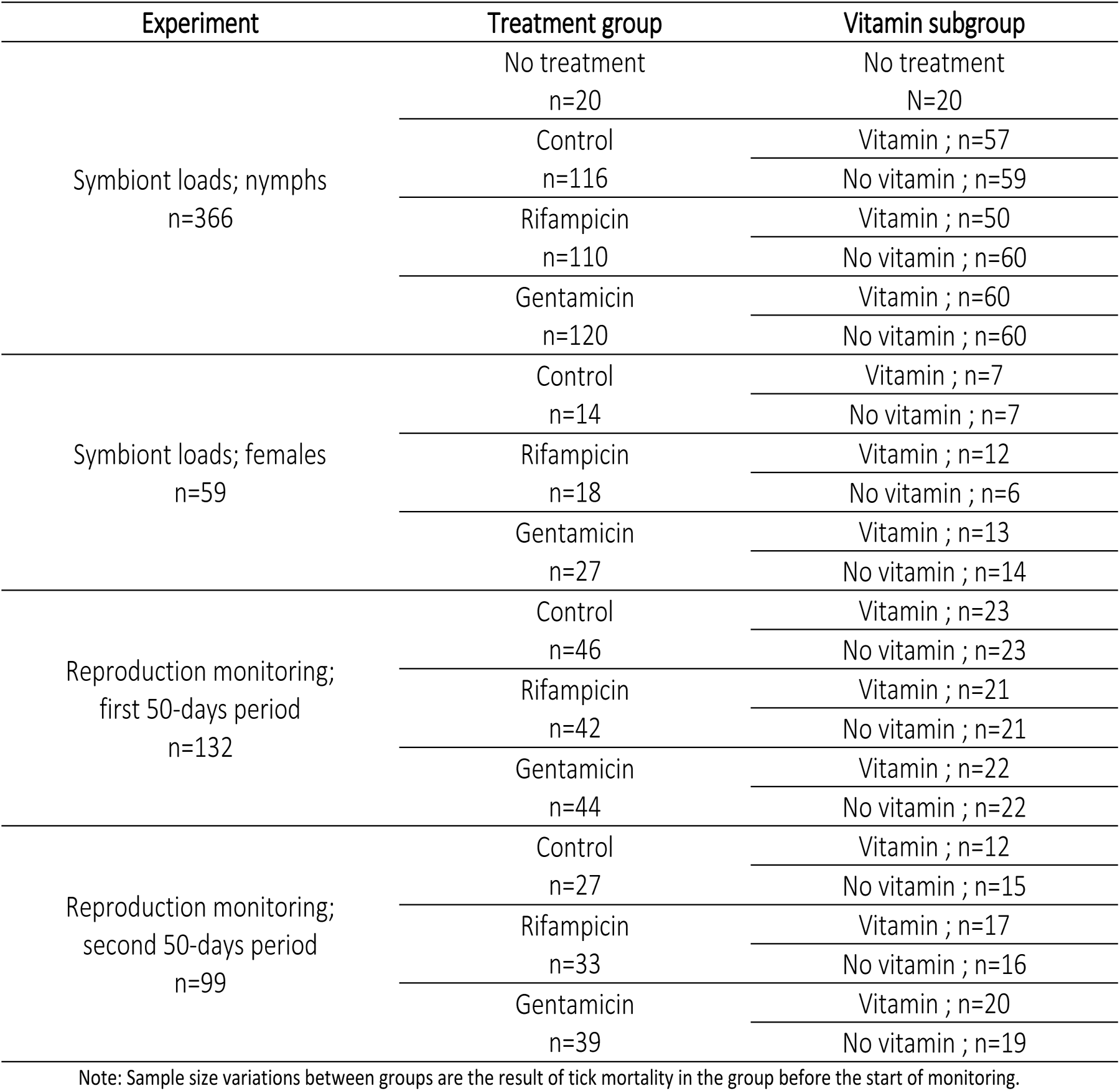
Sample size and repartition of individuals into groups for the different experiments performed.

**Table S2:**
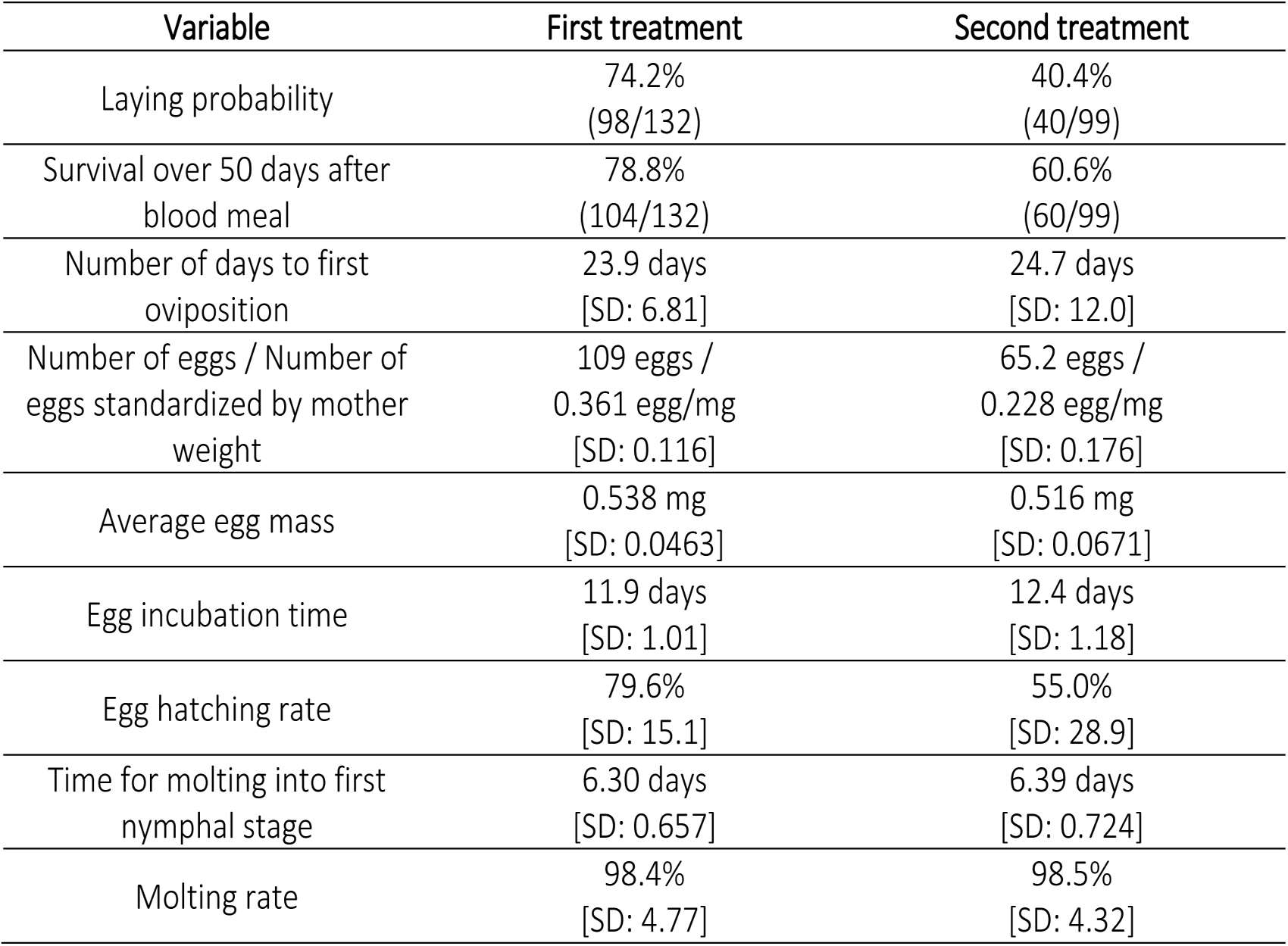
Mean and variance for the traits associated with reproductive fitness after antibiotic treatment and vitamin supplementation.

